## Supplementary Materials for "Deep Generative Models for Discrete Genotype Simulation"

#### 1 Principal Component Analysis (PCA) Results on Genotype Datasets

| Dataset | Number of PCs | MSE | SNP Matching Accuracy |
|---|---|---|---|
| Cow CHR 14 (1,771 SNPs) | 155 | 0.0290 | 97.1089% |
| Cow CHR 5 (2,238 SNPs) | 206 | 0.0281 | 97.1995% |
| Cow All CHRs (50,161 SNPs)* | 4,819 | 0.0277 | 97.2411% |
| Human Ensembl (3,493 SNPs) | 2,026 | 0.0339 | 96.7983% |
| Human CHR 6 (12,283 SNPs) | 5,149 | 0.0143 | 98.5745% |
| Human CHR 12 (9,780 SNPs) | 4,438 | 0.0143 | 98.5726% |
| Human Multi CHRs (42,409 SNPs)* | 18,769 | 0.0144 | 98.5688% |

Table 1: Principal Component Analysis (PCA) performed on each dataset with 90% of the variance retained from the original data. The continuous values are then reconstructed to discrete values (0, 1, or 2) by assigning the closest discrete value, for example, $(-\infty, 0.5) \to 0$, $[0.5, 1.5] \to 1$, and $(1.5, +\infty) \to 2$.

* We apply PCA to each chromosome individually, and then concatenate the results.

### 2 Failed thresholding strategy when attempting to map generated continuous values to 0/1/2

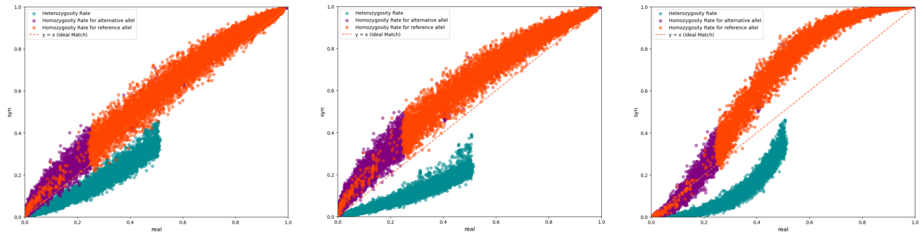

(a) (b) (c)

Figure 1: Comparison of genotype frequencies under different thresholding strategies used to map continuous values to discrete set 0, 1, 2. (a) Thresholds at $[0, \frac{1}{2}, \frac{3}{2}, 2]$, (b) Thresholds at $[0, \frac{2}{3}, \frac{4}{3}, 2]$, and (c) Dynamic thresholding for each SNP based on Hardy–Weinberg equilibrium and allele frequency.

Additionally, it is possible to construct a synthetic population with exactly the same genotype frequencies as the original population. This can be achieved by first ranking all the continuous outputs and then mapping them to genotype categories (0/1/2) according to the frequency proportion observed in the original data. While this approach ensures perfect alignment in genotype frequency, it tends to degrade performance on other metrics. In particular, the recall metric typically drops to 0, indicating poor diversity in the generated samples.

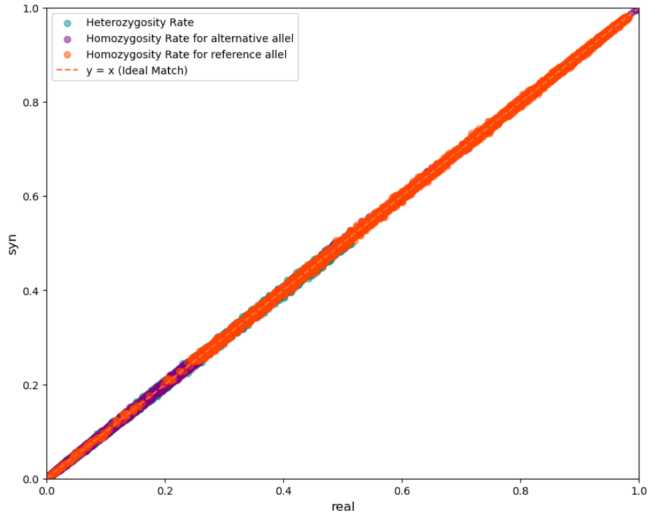


Figure 2: Exact genotype frequency matching between real and synthetic populations using a thresholding strategy based on genotype frequency.

### 3 Precision and Recall with Different Distance Metrics

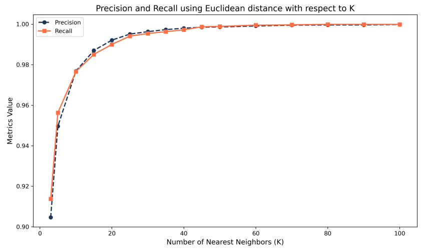


(a) Euclidean Distance on Cow

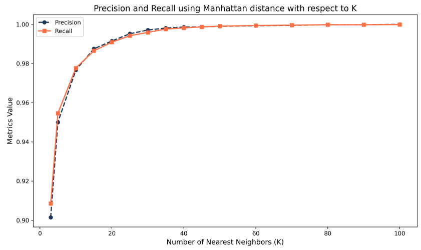


(b) Manhattan distance on Cow

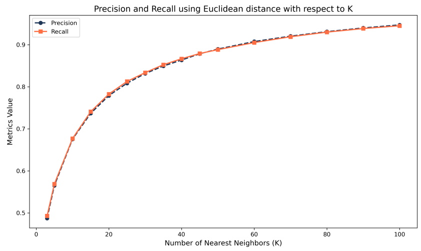


(c) Euclidean Distance on Human

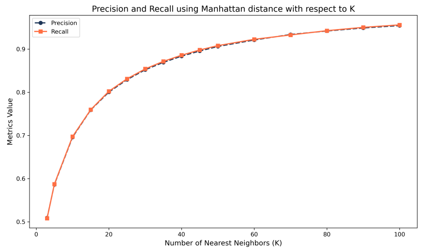


(d) Manhattan Distance on Human

Figure 3: Evolution of Precision and Recall with respect to $k$ using Euclidean and Manhattan Distances on Cow and Human Datasets.

### 4 Method for Calculating Yield Deviation of Fat Content

Fat content is measured by mid-infrared spectrometry on individual milk samples collected during monthly test days. For selection purpose, the trait analyzed is the average value over the entire lactation, with records repeated across lactations. To account for this, we use a multiplicative mixed model that adjusts for heterogeneous phenotypic variances within herd-year and region-year classes and accommodates multiple records per cow. The model accounts for fixed environmental effect, random permanent environmental effect, random genetic effect, and residual effect.

Let $i$ index cows, $j$ index repeated records for each cow, and $h$ index the level corresponding to variance heterogeneity. The $j^{th}$ record for cow $i$, associated with heterogeneity level $h$, is denoted by $y_{i,j,h}$. Formally, it is modeled as:

$$
\begin{aligned}
y_{i,j,h} = (&\text{effect}_{\text{herd}\times\text{year}} \\
&+ \text{effect}_{\text{age at calving}\times\text{region}\times\text{year}} \\
&+ \text{effect}_{\text{calving year}\times\text{calving month}\times\text{region}\times\text{year}} && \text{fixed env effect} \\
&+ \text{effect}_{\text{dry period length}\times\text{region}\times\text{year}} \\
&+ b_i && \text{permanent env effect} \\
&+ a_i && \text{genetic effect} \\
&+ e_i) && \text{residual effect} \\
&\times e^{\gamma_h/2} && \text{heterogeneous variance adjustment factor}
\end{aligned}
$$

where we have:

- **fixed env effect**: fixed effect associated with different combinations of environmental factors.
- **permanent env effect**: random effect accounting for the permanent environmental effect specific to cow $i$.
- **genetic effect**: additive genetic effect of cow $i$ such that $\text{Var}(a) = \mathbf{G}\sigma_a^2$, where $\mathbf{G}$ is the genomic relationship matrix and $\sigma_a^2$ is the additive genetic variance.
- **residual effect**: random residual error term for cow $i$ such that $\text{Var}(e) = \mathbf{I}\sigma_e^2$, where $\sigma_e^2$ is the residual variance.
- **heterogeneous variance adjustment factor**: $\gamma_h$ is the parameter modeling heterogeneity in phenotypic variance, decided by [herd $\times$ year] and [region $\times$ year] [1]. The factor $e^{\gamma_h/2}$ is applied to standardize variance across environments.

In our study, the estimated additive genetic variance $\sigma_a^2$ is 8.84, the variance of the permanent environmental effect is 3.54, and the residual variance $\sigma_e^2$ is 5.3.

Therefore, the heritability $h^2$ of the fat content trait is $8.84/(8.84+3.54+5.3) = 0.5$.

We first adjust each performance record $j$ of cow $i$ to account for heterogeneous variance, fixed environmental effects, and the permanent environmental effect. The adjusted record, denoted $y_{i,j}^{\text{adjusted}}$, is given by

$$y_{i,j}^{\text{adjusted}} = y_{i,j,h} \times e^{-\gamma_h/2} - \text{fixed env effect} - \text{permanent env effect}$$

Finally, the yield deviation $YD_i$ for cow $i$ is computed as a weighted average of all its adjusted records, given by

$$YD_i = \frac{\sum_{j=1}^{3} w_{i,j} \times y_{i,j}^{\text{adjusted}}}{\sum_{j=1}^{3} w_{i,j}},$$

where $w_{i,j}$ reflects the amount of information contained in the yield deviation, accounting for the number of elementary records per cow, the size of the environmental effect groups, the repeatability, and the heritability of the trait.

### 5 Neural Network Architecture and Hyperparameter Selection for All Chromosomes of the Cow Dataset

#### 5.1 VAE

**Model Architecture**

- **Input dimension:** 150,483 features, corresponding to one-hot encoded genotypes for 50,161 SNPs.
- **Encoder:**
  - Fully connected layers: 150,483 → 4,096 → 2,048 → 1,024
  - One residual block at 1,024 units (2 linear layers with skip connection)
  - Linear projection to 512, followed by two separate linear heads to infer:
    - Mean vector $\mu \in \mathbf{R}^{256}$
    - Log-variance vector $\log \sigma^2 \in \mathbf{R}^{256}$
  - All layers are followed by Batch Normalization to stabilize training and LeakyReLU activation (slope = 0.05) to avoid neuron inactivation.
- **Latent space:** 256-dimensional latent vector
- **Decoder:**

  - Fully connected layers: 256 → 512 → 1,024
  - One residual block at 1,024 units (same structure as encoder)
  - Continuation: 1,024 → 2,048 → 4,096 → 150,483 (The reverse one-hot decoding is handled during the loss computation, not as an explicit output layer)
  - All hidden layers include Batch Normalization and LeakyReLU (slope = 0.05)

**Training Hyperparameters**

- **Batch size:** 2,048
- **Optimizer:** Adam optimizer with $\beta_1 = 0.9$, $\beta_2 = 0.999$
- **Learning rate:** $5 \times 10^{-4}$

##### 5.2 GAN

**Model Architecture**

- **Generator**
  - **Input:** 256-dimensional latent vector
  - **Initial transformation:**
    * Linear layer: 256 → 1,045
    * Batch Normalization + LeakyReLU
  - **Residual blocks:** Two blocks with intermediate expansion:
    * Block 1: 1,045 → 1,045 (residual) → 2,090
    * Block 2: 2,090 → 2,090 (residual) → 4,180
    * Each residual block consists of two fully connected layers with skip connection, Batch Normalization, and LeakyReLU
  - **Final projection:**
    * Linear: 4,180 → 150,483 (matches the one-hot encoded genotype size)
    * Reshape to [batch size, 50161, 3]
    * Apply Gumbel-Softmax to obtain discrete-like output while maintaining differentiability
- **Discriminator**
  - **Input:** 150,483-dimensional vector
  - **Initial transformation:**
    * Linear layer: 150,483 → 4,180
    * LeakyReLU activation

  - **Residual blocks:** Two blocks with progressive contraction:
    * Block 1: 4,180 → 4,180 (residual) → 2,090
    * Block 2: 2,090 → 2,090 (residual) → 1,045
    * Each residual block includes two linear layers and LeakyReLU activations
  - **Final layer:**
    * Linear: 1,045 → 1
    * Sigmoid activation for binary classification.

- **Activation functions:** All hidden layers use LeakyReLU (negative slope = 0.05)

- **Normalization:** Batch Normalization is used in the generator but omitted in the discriminator

**Training Hyperparameters**

- **Batch size:** 128

- **Optimizer:** Adam optimizer with $\beta_1 = 0.5$, $\beta_2 = 0.999$ for both generator and discriminator

- **Learning rate:** $1 \times 10^{-4}$ for both generator and discriminator

- **Gumbel-Softmax temperature:** Linearly annealed from 1.0 to 0.1 during training

#### 5.3 WGAN

Here we present the conditional version, where phenotype information is provided during training. For the unconditional setting, it suffices to remove the phenotype-related tensors and adjust the tensor shapes accordingly.

**Model Architecture**

- **Generator:**
  - **Input:** A 260-dimensional vector composed of:
    * Latent noise vector $z$ of dimension 256
    * Phenotype conditioning vector, repeated and concatenated of dimension 4
  - **Initial Layer:**
    * Linear: 260 → 1,045
    * Batch Normalization + LeakyReLU
  - **Residual Blocks:** Two sequential residual blocks:

    * **Block 1:**
        · ResNetBlock: A phenotype vector of dimension 4 is first injected, resulting in an input dimension of 1,049. The block consists of two linear layers with Batch Normalization and a skip connection, maintaining the dimensionality at 1,049.
        · Linear: 1,049 → 2,090 + LeakyReLU
    * **Block 2:**
        · ResNetBlock: Similarly, a phenotype vector of dimension 4 is injected, resulting in a dimension of 2,094. The block maintains this dimensionality through two linear layers with Batch Normalization and a skip connection.
        · Linear: 2,094 → 4,180 + LeakyReLU
  - **Final projection:**
    * Linear: inject phenotype vector of dimension 4 → 4,184 → 150,483
    * Reshape to [batch size, 50161, 3]
    * Apply Gumbel-Softmax

- **Critic:**
  - **Input:** A flattened genotype sequence of dimension 150,483 is concatenated with a phenotype vector of dimension 4, resulting in a total input dimension of 150,487.
  - **Initial Layer:**
    * Linear: 150,487 → 4,180
    * LeakyReLU
  - **Residual Blocks:**
    * **Block 1:**
        · ResNetBlock: inject phenotype vector of dimension 4 → 4,184 → 4,184
        · Linear: 4,184 → 2,090 + LeakyReLU
    * **Block 2:**
        · ResNetBlock: inject phenotype vector of dimension 4 → 2,094 → 2,094
        · Linear: 2,094 → 1,045 + LeakyReLU
  - **Final output:**
    * Linear: 1,045 → 1 (critic score)

- **Activation functions:** All hidden layers use LeakyReLU (negative slope = 0.05)

- **Normalization:** Batch Normalization is used in the generator but omitted in the critic

##### Training Hyperparameters

- **Batch size:** 128
- **Optimizer:** Adam optimizer with $\beta_1 = 0.5$, $\beta_2 = 0.9$ for both generator and critic
- **Learning rate:** $1 \times 10^{-4}$ for both generator and critic
- **Critic updates per generator update:** 5
- **Gradient penalty coefficient ($\lambda$):** 10
- **Gumbel-Softmax temperature:** Linearly annealed from 1.0 to 0.1 during training

## 5.4 DM

Similarly, we provide the conditional version here. To switch to the unconditional setting, simply remove all phenotype-related tensors and adjust the tensor shapes accordingly.

##### Model Architecture

- **Input structure:**
  - The input tensor is a concatenation of:
    - Noisy data vector $x \in \mathbf{R}^{4819}$ (corresponding to the PCA-latent genotype data)
    - Time embedding $t_{\text{emb}} \in \mathbf{R}^{256}$
    - Phenotype embedding $pheno_{\text{emb}} \in \mathbf{R}^{64}$
  - Total input dimension: 5,139
- **Time embedding:**
  - Uses a sinusoidal positional encoding, similar to that in the original DDPM implementation.
  - Embedding dimension is set to 256
- **Phenotype embedding:**
  - A continuous label (scalar) is projected to a higher-dimensional space using a linear layer
  - Embedding dimension is set to 64
- **Noise predictor architecture:**
  - `fc1`: Input layer mapping from 5,139 $\rightarrow$ 8,192

- After the first hidden layer, time and phenotype embeddings are reinjected (8192 + 256 + 64 = 8512)
- `fc2`: 8,512 → 8,192
- `fc3`: 8,192 → 6,144
- `out`: Final output layer from 6,144 → 4,819
- **Residual connection:**
  - `res`: A skip connection projects the input via a linear layer: 5,139 → 4,819
  - The final output is computed as `out + res`
- **Normalization:** Layer Normalization is applied after each internal fully connected layer
- **Activation:** ReLU is used after each normalization layer

**Training Hyperparameters**

- **Batch size:** 4,086
- **Diffusion process:**
  - Number of diffusion steps: 1,500
  - $\beta$ schedule: linear
- **Time Sampling Strategy:** Antithetic sampling
- **Optimizer:**
  - Adam optimizer with $\beta_1 = 0.9$, $\beta_2 = 0.999$
  - Learning rate: $3 \times 10^{-4}$
  - Learning rate scheduler: Cosine Annealing
  - Minimum learning rate: $3 \times 10^{-6}$
  - Warm-up:
    - Strategy: Linear warm-up
    - Period: first 1,000 steps

### 6 Impact of SNP Dependence on the Difficulty of Generative Modeling

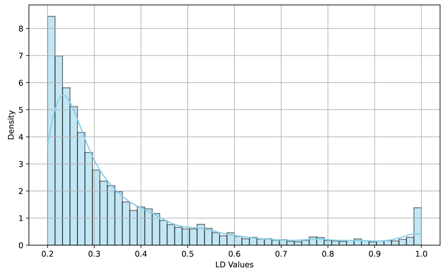


(a) Cow CHR 5

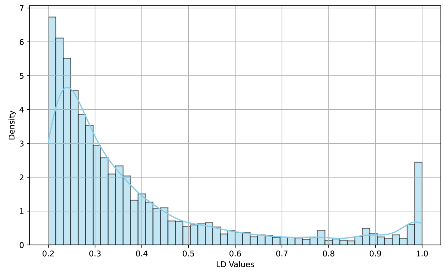


(b) Cow CHR 14

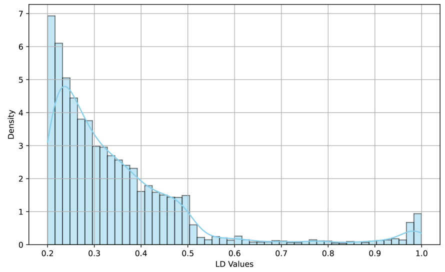


(c) Human CHR 6

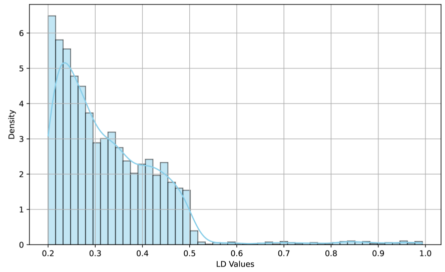


(d) Human CHR 12

Figure 4: Distribution of LD values across different datasets, focusing on values $\geq 0.20$. In Cow dataset (a and b), SNPs exhibit stronger correlations compared to the Human dataset (c and d). In Human dataset, Chromosome 6 (c) contains more high-LD SNP pairs than Chromosome 12 (d). This makes it easier for generative models to learn Chromosome 6, despite its larger dimension compared to Chromosome 12.

### 7 Limitation of AA Score as a Robust Metric

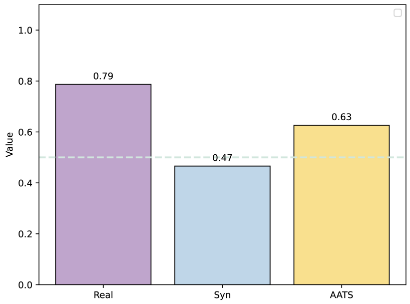


(a) WGAN on Cow CHR 5

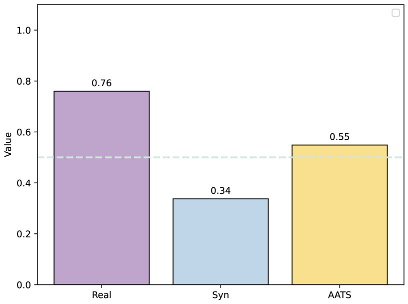


(b) WGAN on Cow CHR 14

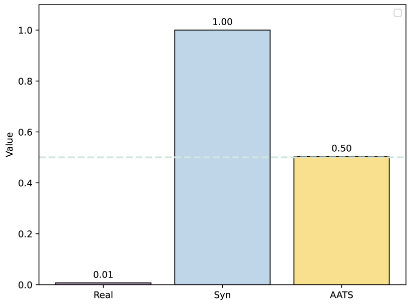


(c) DM on Human CHR 6

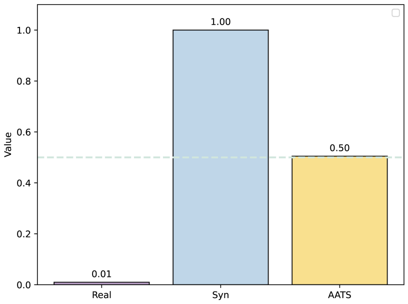


(d) DM on Human CHR 12

Figure 5: A detailed investigation of the $AA$ score: (a) and (b) show cases where the $AA$ score serves as a good evaluation metric, with both $AA_{real}$ and $AA_{syn}$ yielding good scores, resulting in an $AA$ score around 0.50. (c) and (d) depict an anomalous scenario where $AA_{real} \approx 0$ and $AA_{syn} \approx 1$, yet the $AA$ score remains around 0.50, highlighting a limitation of using $AA$ as a metric.

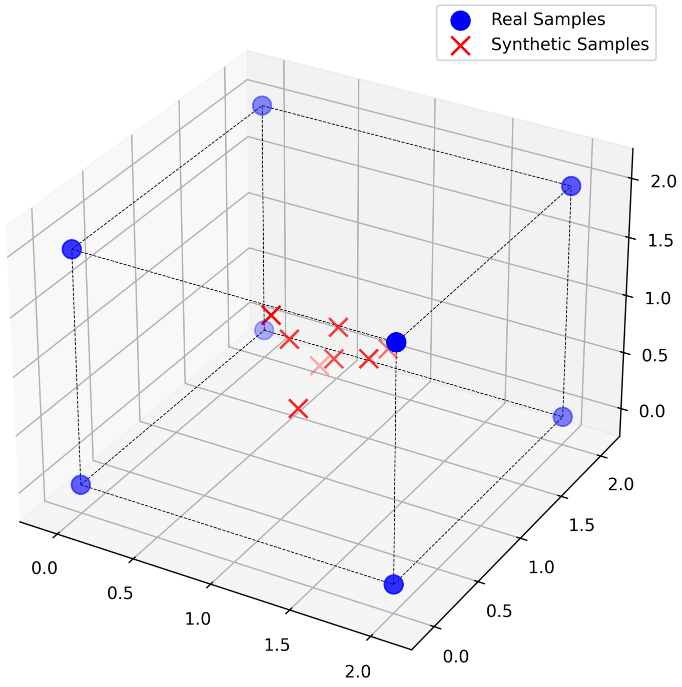


Figure 6: Geometric illustration of a scenario where $AA_{real} = 0$ and $AA_{syn} = 1$: Eight real samples are positioned at the corners of a cube, with each pair of real samples' distance equal to 2 ($d_{RR} = 2$). Eight synthetic samples are tightly clustered at the center of the cube [1, 1, 1]. In this setup, each real sample is closer to a synthetic sample than to any other real samples ($d_{RS} < \sqrt{3} < d_{RR}$), yielding $AA_{real} = 0$. Conversely, each synthetic sample is closest to another synthetic sample in the cluster, yielding $AA_{syn} = 1$. If we relate this scenario to the precision and recall metrics with $\forall k$, we obtain a precision of 1 because every synthetic sample falls within the support of a real sample. However, the recall is 0, as none of the real samples fall within the support of any synthetic sample.
